# Female Excellence in Rock Climbing Likely Has an Evolutionary Origin

**DOI:** 10.1101/2020.05.26.116244

**Authors:** Collin Carroll

## Abstract

The human body is exceptional for many reasons, not the least of which is the wide variety of movements it is capable of executing. Because our species is able to execute so many discrete activities, researchers often disagree on which were the movements most essential to the evolution of our species. This paper continues a recently introduced analysis, that the performance gap between female and male athletes narrows in sports which most reflect movements humans evolved to do. Here, I examine the performance gap in rock climbing. Because rock climbing is so similar to tree climbing, which bountiful evidence suggests has been key to the origin and proliferation of our species, we would expect to see a narrow performance gap between men and women in the sport. Indeed, this is the case. Female climbers are some of the best in the world irrespective of gender, a trend that is not found in any other major sport. I conclude that exceptional ability of female climbers is further evidence of the existence of sex-blind musculoskeletal adaptations, which developed over the course of human evolution to facilitate essential movements. These adaptations abate the general physical sexual dimorphism which exists in humans. This paper provides more evidence that the performance gap in sport can be used as a measure of human evolution.

## Introduction

While evolutionary anthropologists generally agree on much of the history of human development, there is still a great deal of debate as to which movements the human body has adapted. Most of this debate is specific to those movements that arose in the past few million years in conjunction with the genus *Homo*. Some anthropologists, for instance, claim that the ecological dominance of humans can be primarily attributed to our overhand throwing ability (Lombardo and Deaner 2018a; Roach, Venkadesan, Rainbow, & Lieberman 2013; Wilson, Zhu, Barham, Stanistreet, & Bingham 2016), another sect argues that endurance running has been critical to the development of our species (Bramble and Lieberman 2004; Liebenberg 2006; Lieberman, Raichlen, Pontzer, Bramble, & Cutright-Smith 2006), and others still make the case that humans adapted to excel at intraspecies hand-to-hand combat (Carrier 2011; Carrier, Schilling, & Anders 2015). These are only a few perspectives on the evolutionary history of *Homo sapiens*, each supported with its own set of evidence.

It is generally accepted, however, that for the millions of years before the emergence of *Homo*, human ancestors were capable tree climbers (Green and Alemseged 2012; Stern and Susman 1983; Wood and Baker 2011). Even now, as is the case with other primates, humans remain capable climbers; a number of researchers have observed modern hunter-gatherers climbing trees to acquire resources (Ichikawa 1981; Venkataraman, Kraft, & Dominy 2012). Arboreal locomotion was an essential part of our development as a species.

The importance of climbing is imprinted in the human musculoskeletal system: our long arms, short trunk, and upright posture all appear to have originated to facilitate arboreal locomotion (Crompton, Vereecke, & Thorpe 2008; Fleagle 2013; Thorpe, Holder, & Crompton 2007). These physiological features are seen in both the male and female members of our species, and with that in mind, this paper aims to build on a recently introduced idea: that a narrower performance gap (PG) between males and females has an evolutionary cause (Carroll 2019). In other words, sports that show a smaller difference in ability between male and female athletes are those most similar to movements essential to the evolution of *Homo sapiens*. This finding would be notable because it would add another metric to evaluate the suitability of the human body to certain movements, and it could help refine the narrative of human evolution.

Climbing, with the deep reservoir of archeological and physiological evidence that suggests its importance to the development of *Homo sapiens*, would therefore be expected to have a particularly narrow PG. This paper examines the PG between males and females in rock climbing, the sport most similar to the arboreal movement that was essential to the origination of our species, to identify whether climbing does indeed have a narrow PG. If true, this would provide more evidence that the PG is a useful tool for measuring the evolutionary relevance of certain movements.

### Evolutionary Evidence of Arboreal Movement to the Development of Modern Humans

There is considerable evidence supporting the idea that arboreal movement was crucial to the survival of early humans. Before the emergence of the genus *Homo*, human ancestors relied on a lifestyle that seems to be a mix of arboreal and terrestrial in nature to avoid predators and access resources (Senut 2014; Stern 2000). And while tree-climbing behavior in early humans almost certainly preceded bipedalism, there is debate as to whether bipedalism first descended from upright posture in the trees or whether it emerged from knuckle-walking patterns of terrestrial locomotion (Begun 2004; Gebo 1996; Prost 1980; Schmitt 2003; Senut, Pickford, Gommery, & Ségalen 2018; Stern and Susman 1981). Regardless of the exact transition from arboreal to terrestrial locomotion in humans, it is clear our ape ancestors led a life in the trees.

While the exact degree to which *Australopithecus afarensis* moved and lived arboreally is in dispute, it seems clear that the species spent a significant amount of time in the trees and held adaptations needed for arboreal locomotion (Crompton, Vereecke, & Thorpe 2008; Crompton et al. 2011; Duncan, Kappelman, & Shapiro 1994; Green and Alemseged 2012; Stern 2000). The fossil record of the australopith A.L. 288-1 (“Lucy”), for instance, shows evidence of substantial reliance on climbing behavior (Jungers 1982; Kappelman et al. 2016; Meyer, Williams, Smith, & Sawyer 2015; Ruff, Burgess, Ketcham, & Kappelman 2016).

The importance of tree climbing is not limited to our distant ancestors, however; evidence of arboreal locomotion can be seen in the genus *Homo*, too. Antón and Snodgrass (2012), comparing *Homo* to *Australopithecus*, found that both genera have a locomotor repertoire that is significantly dependent on arboreal movement. Indeed, early hominins do evince a number of features that can be linked to frequent arboreal behavior (Roberts, Boivin, Lee-Thorp, Petraglia, & Stock 2016; Susman, Stern, & Jungers 1984; Tuttle 1981). One of these features, for instance, is a robust upper limb that suggests a high tolerance for mechanical loading, which would have been necessary for a life spent at least partially in trees (Haeusler and McHenry 2007; Heinrich, Rose, Leakey, & Walker 1993; Ruff 2009; Susman and Creel 1979). Features like this provide evidence of the importance of arboreal locomotion to early *Homo*.

Beyond the importance it had to early *Homo*, arboreal locomotion appears to be significant to our own species, too. Of course, *Homo sapiens* is better suited to bipedal terrestrial locomotion than to arboreal locomotion, given how energetically economical walking is compared to climbing in humans (Elton, Foley, & Ulijaszek 1998; Kozma et al. 2018). But even with this clear preference, humans do retain many traits that facilitate tree climbing. Compared to smaller, primarily arboreal primates, humans do not expend significantly more energy climbing per kilogram of body mass, meaning we are still efficient climbers (Hanna, Schmitt, & Griffin 2008). The human hand, with its “power” grip designed to facilitate prehensile movements as described by Napier (1956), also allows for arboreal locomotion; additionally, humans have been measured using subtle proprioceptive measures, like “light touch” fingertip support, to reduce bipedal instability and improve balance in tree-canopy-like environments (Johannsen et al. 2017). Even the gluteus maximus muscle, essential for bipedal locomotion, is particularly active during climbing, suggesting that some adaptations which favor terrestrial bipedalism also have use in arboreal movement (Bartlett, Sumner, Ellis, & Kram 2013). The human mind also appears to have arboreal origins as well. Povinelli and Cant (1995) suggest the idea of self-conception – the cognitively advanced ability that allows an organism to perceive of itself – in large great apes and humans originally developed to better facilitate arboreal clambering. And an experiment of human tree climbers by Hanson (2016) found that most made conscious choices to better facilitate their movement up a tree. Perhaps the most compelling evidence of human tree climbing, though, comes from behavioral studies of select modern hunter-gatherer populations, who frequently climb trees as a method of resource acquisition (Kraft, Venkataraman, & Dominy 2014). In short, while the exact degree to which tree climbing has been useful to *Homo sapiens* is yet undetermined, plenty of evidence suggests that humans remain well adapted to arboreal locomotion.

### Rock Climbing as a Modern Analog to Tree Climbing

As a result of millions of years of arboreal locomotion, the ability to climb is ingrained in the human form. This ability is not limited to climbing trees exclusively; certain physiological and cognitive traits found in humans also facilitate rock climbing. One is the strength of the human hand. Grip and finger strength are unsurprisingly crucial in rock climbing; elite rock climbers possess stronger grips and better finger and hand endurance than less experienced rock climbers and non-climbers (Cutts and Bollen 1993; Ozimek et el. 2017; Philippe, Wegst, Müller, Raschner, and Burtsche 2011). Additionally, the position of the human body during climbing further improves grip strength. According to Parvatikar and Mukkannavar (2009), grip strength is highest when the shoulder is positioned in 180° of flexion, the exact positioning of a climber grasping onto a hold above his or her body. Our muscular endurance is another key to our species’ rock-climbing ability; humans appear to have a greater relative VO_2_max – a measurement of an organism’s maximal oxygen consumption – than other apes (Pontzer 2017). Booth, Marino, Hill, & Gwinn (1999) suggest that “outdoor rock climbing might require a large fraction of the climber’s peak oxygen uptake,” so our high VO_2_max advantages climbing in humans. The human mind is crucial to our rock-climbing ability, too. Rock climbers, when compared to untrained controls, have improved visual-spatial perception (Marczak, Ginszt, Gawda, Berger, & Majcher 2018), and expert rock climbers also have better visual and motor memory when compared to climbers with less expertise (Whitaker, Pointon, Tarampi, & Rand 2019). Because rock climbing and tree climbing are similar, and because competition tree climbing is effectively nonexistent, rock climbing serves as the athletic analog of tree climbing in this paper.

### The Performance Gap in Sport as a Measure of Human Evolution

This paper expands on the idea that the PG between across sports shrinks or narrows depending on how relevant a sport is to human evolution. In recent years, the PG on the whole has stabilized, meaning the current difference between male and female athletes at any given sport is a good indicator for the difference between overall male and female ability – the idea being that elite athletes reflect the upper limits of human capability (Millard-Stafford, Swanson, & Wittbrodt 2018). The previous paper I published on the subject of the PG as a tool for measuring evolutionary movements dealt exclusively with track & field events; the extremely high-quality data from that study strongly suggested that there was a significantly narrower PG in short-distance sprinting events compared to longer-distance events, and that there was also a significantly narrower PG in running events of all distances compared to jumping events (Carroll 2019). Because it was evolutionary advantageous for humans, regardless of gender, to sprint well – to escape predators, primarily – I argue that this selection pressure led to an accrual of traits that made for better sprinters in both males and females. I termed these traits sex-blind musculoskeletal adaptations (SBMA’s). SBMA’s bridge the general gap of physical dimorphism that exists between men and women, which in turn leads to a narrower PG in the sports most similar to movements humans adapted to do.

Because it is analogous to the arboreal locomotion instrumental to the development of our species, then, the sport of rock climbing would be expected to have a particularly narrow PG. The purpose of this paper is to explore whether or not this hypothesis holds true. If it does, this paper would further provide further evidence that a narrow PG between men and women in a sport stems from the heavy reliance early humans had on comparable movements. The PG could then be used as a metric to gauge how well-equipped the human body is to certain activities – as a result of accrued SBMA’s – and could help clarify the evolutionary history of *Homo sapiens*.

## Materials and Methods

### How Climbing Ability is Assessed

Unlike many single-competitor sports such as track & field or Olympic lifting, which are standardized to allow for easy comparison between competitors, ability in rock climbing is measured by the competitors, who “rate” the climbs they complete based on difficulty. In this manner, the athletes in rock climbing act as the judges of the sport as well as its record keepers. Once a rock climber “sends,” or completes, a particular route for the first time, he or she is responsible for its rating so that other climbers will know the difficulty of the route. These athletes grade rock climbing routes according to different scales; for the sake of clarity, the Yosemite Decimal System (YDS) is used in this paper unless otherwise noted.

The YDS is divided into 5 main classes, with Class 5 used to grade rock climbing routes. Class 5 is divided into categories, with 5.15 being the most difficult that currently exists. Further subcategories a-d exist within each route categorized as 5.10 and above. Thus “Silence,” the most difficult rock-climbing route ever sent, holds a rating of 5.15d (Skenazy 2017). Climbing ability tends to be measured as a climber’s “best ascent” – that is, the most challenging route a climber has sent – to measure his or her skill in the sport. Only those elite climbers who have sent a 5.15-rated route are noted here.

### Climbing Data

This paper uses data aggregated by rock climber William Kuelthau published on the website Rock & Ice (2020). This list was last updated on February 4^th^, 2020, and the data gathered and analyzed on March 24, 2020. The climbers who have sent a 5.15-grade route, 90 in all, are included in this data set by best ascent so that no climber is recorded more than once. The data here are measured conservatively, so any climb specifically marked as questionable by Kuelthau will not be included in the set.

### Limitations of Climbing Data

The data on outdoor rock climbing suffers chiefly from a lack of standardization, bifurcated into two main issues: first, rock climbing records are not held by any dedicated organization; second, there is some subjectivity in the way routes are rated by climbers.

Rock climbing as a sport lacks an official record keeper responsible for tracking which athletes have sent which routes. Organizations like the International Federation of Sport Climbing exist, but their purpose is more to establish rock climbing competitions on artificial courses than to keep track of outdoor free climbing and bouldering ascents. By contrast, track & field is governed by an official body, World Athletics, which maintains records for all official track & field events for both men and women (“World Athletics” 2020). The lack of an official body governing outdoor rock climbing means the data on ascents is of inherently lower quality than that of the data on track & field events. Because climbing results are self-reported rather than recorded at official events, unscrupulous climbers have more freedom than track athletes to make dishonest claims about whether they have actually completed the routes which they have claimed. However, this is unlikely to significantly affect climbing results at the highest level, 5.15 and above, because the attempts of these climbers are typically witnessed by others and the results often recorded, providing a record of the ascents (“Gripped” 2017). As stated earlier, any climbs marked as questionable are not included in my data set.

Another issue with rock climbing records is that they are measured qualitatively, by the climbers who are the first to send each route. Track performance, on the other hand, is measured quantitatively, by an athlete’s time to run a certain distance, for instance. The subjectivity of rock-climbing records, while problematic, is however mitigated to an extent. Experienced rock climbers do indeed tend to rate routes accurately, according to a study by Draper et al. (2011). If the original climber gives a route an inappropriate rating – one that is too easy or too difficult for the route – it can be changed based on the collective input of subsequent climbers (Pesterfield 2018). This dynamism does mitigate some of the subjectivity of the rating system, especially at the upper reaches of the sport, where rating difficult routes can become a collaborative effort between the few climbers capable of ascending the world’s toughest climbs (Lucas 2017). Because ascending a route rated 5.15a or higher is such a rarity in climbing, the added scrutiny makes it more difficult for a climber to inflate an easier route to a grade in the 5.15 range. Fewer than 100 climbers have ever sent a 5.15-rated route, and the most accomplished of these elite climbers – those who have sent a 5.15b or higher, for instance – can confirm or deny 5.15 status for many of these extremely difficult climbs (“Gripped” 2018). For the purposes of this paper, it can be inferred that the ratings of the most difficult outdoor rock climbing routes in the world – 5.15a through 5.15d – tend to accurately reflect the actual difficulty of these routes; any climber who has sent a 5.15-rated route is very likely to be one of the top rock climbers of all time.

## Results

Table 1 and Figure 1 summarize the number of climbers, by gender, who have sent each of the four most difficult ratings, 5.15d through 5.15a. Supplementary data of each of the climbers by name can be found in Appendix 1. With one female climber who has sent a 5.15b-rated route and two others who have sent 5.15a-rated routes, females compose 3 of the top 90 climbers of all time. Figure 2 shows how many male athletes, at minimum, have eclipsed the top female athlete at three different sports: rock climbing, the 100-meter dash, and the marathon. Data on the 100-meter dash and the marathon come from WorldAthletics.com. Because the data listed by World Athletics is cut off at a certain time – 10.30 seconds for the 100-meter dash and 2:12:00 for the marathon – they do not represent the total numbers of male athletes who have eclipsed the top female athlete at each of these two sports.

**Table 1:**
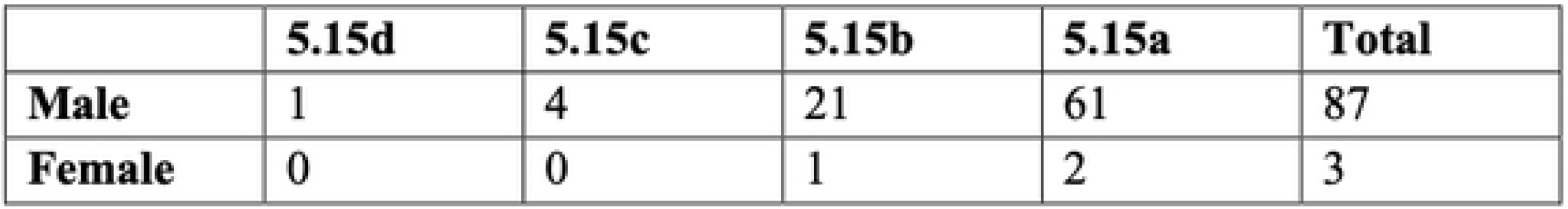
Number of climbers, by gender, who have sent each of the four routes graded most difficult. Climbers are listed once, by their best ascent.

**Figure 1:**
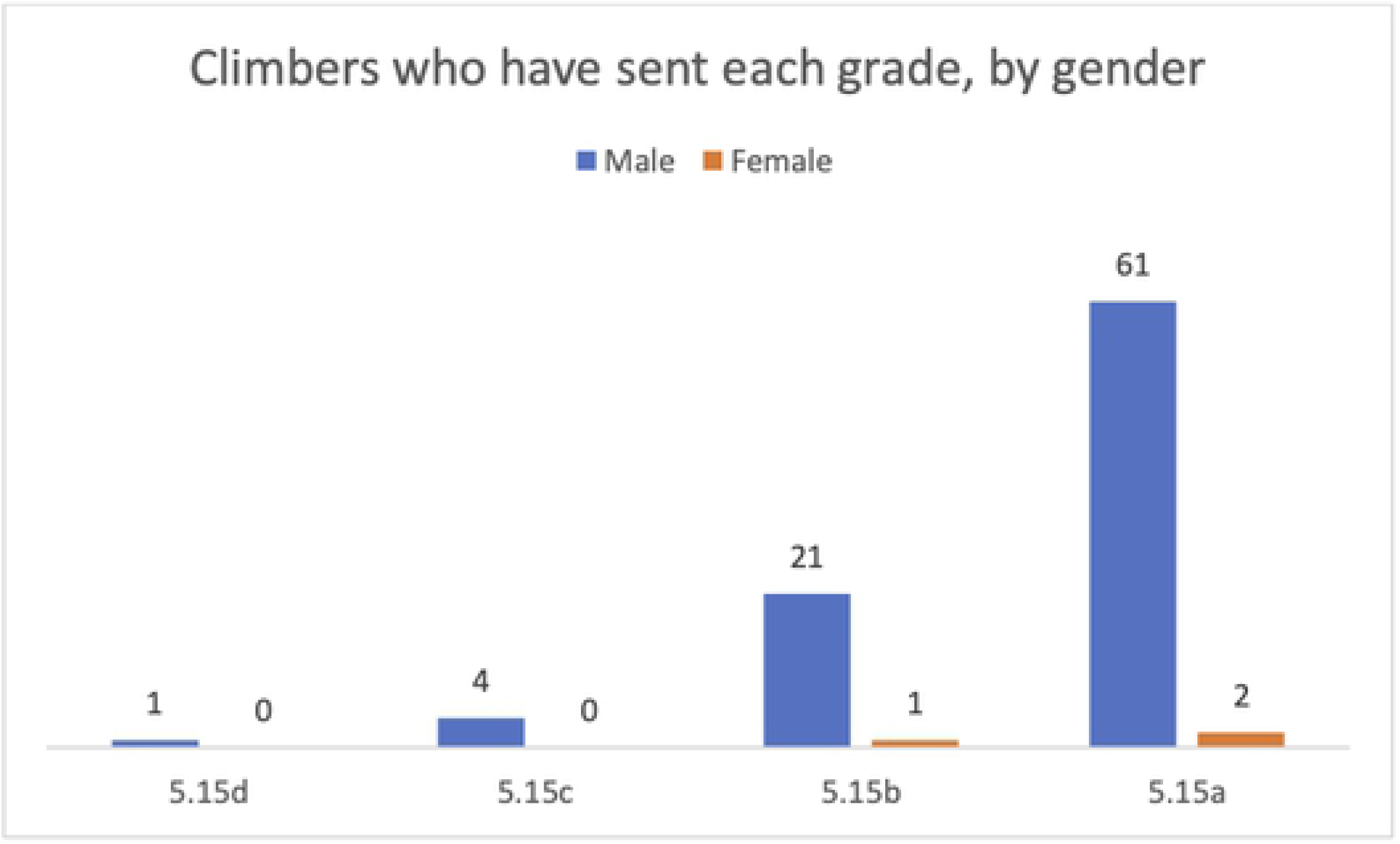
Number of climbers, by gender, who have sent each of the four routes graded most difficult.

**Figure 2:**
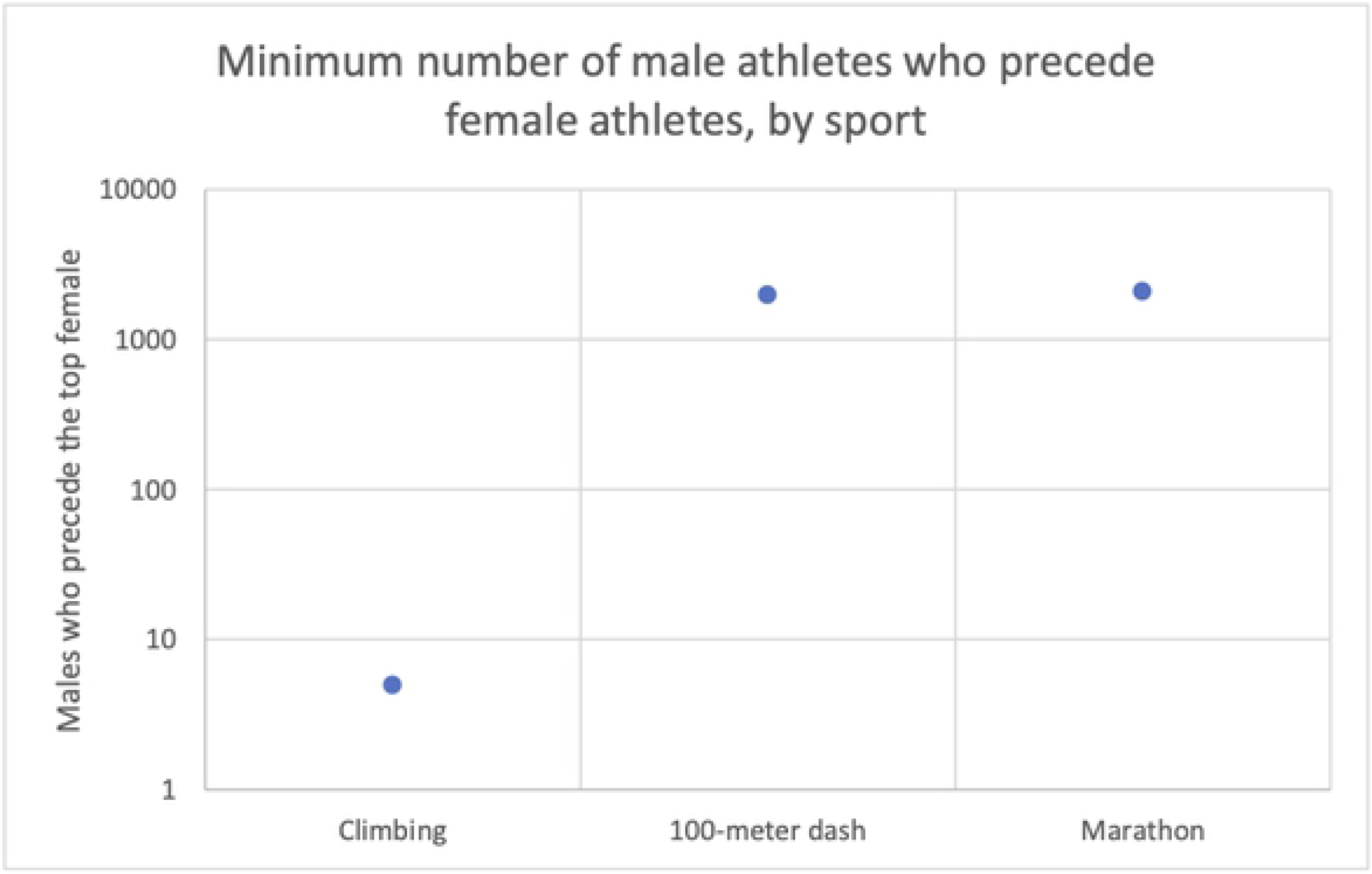
Minimum number of male athletes who have outperformed the top all-time female athlete at three sports. Many more male are likely to precede the top female in both the 100-meter dash and the marathon, but records are limited in each sport.

## Discussion

### Female Ability in Rock Climbing Relative to Other Sports

As hypothesized, relative female rock-climbing ability was shown to be extraordinary. A female climber has sent a 5.15b-rated route, one of only 27 people to have successfully ascended a route rated 5.15b or higher. Additionally, this exceptional performance is not limited to one female outlier. Two other women have sent 5.15a-rated routes, placing 3 females in the top 90 climbers of all time. This level of female achievement is far beyond that seen in other sports. The meter dash, with a relatively narrow PG itself, does not have a single female runner in the top 2,000 competitors, and the fastest female time ever recorded is slower by 0.19 seconds than the fastest-ever male time. This trend holds true for the marathon, too. The top female does not enter into the top 2,000 marathon runners, and she is more than two minutes slower than the 2,000^th^-fastest male. This means that there are likely many thousands more male runners who surpass the world-record-holding females in each event. Rock climbing as a sport shows a much narrower PG at its upper echelon than either short-distance or long-distance running.

This fact provides more evidence that the PG in sport can be used as a metric of human evolution. A sport like rock climbing, with its numerous exceptional female athletes, exhibits a remarkably narrow PG. This is a result of the duration of time early humans spent climbing trees as a means of survival. Arboreal locomotion was essential to our development as a species, and its importance has led to the development of many climbing specific SBMA’s. The idea behind SBMA’s is that certain traits – like a high strength-to-weight ratio, in the case of tree climbing – have been particularly selected for in early humans. These traits bridge the gap of general sexual dimorphism that exists between men and women. Males, due to their body composition and size, hold a general advantage over females in physical activity, but the presence of a greater number of climbing specific SBMA’s shrinks this gap and therefore the PG in climbing sports. Capable tree climbers, male and female, could avoid predators and access resources unavailable to others, allowing them to survive and reproduce.

### Limitations

This analysis of the PG has its limitations. As stated earlier, the data on climbing is not standardized, nor was it able to be gathered from a dedicated rock-climbing organization. Additionally, because the track data offered by World Athletics is cut off at certain times for men, I was unable to accurately count the number of men who out-achieved the overall top woman in both the 100-meter dash or the marathon. Still, the data does appear to accurately reflect how relatively narrow the PG is between the top men and women in rock climbing relative to other sports. Even though the quality of the data may be low, it is still clear that, when compared to other physical activities, women are exceptional rock climbers.

The ratio of male to female rock climbers may also impact the size of the PG; if a greater proportion of athletes at any given sport are women, the PG should be narrower for that sport. The ratio of rock climbers by gender, according to an estimate by professional climber Sasha DiGiulian, is 60% men to 40% women (Dwyer 2019). This would be a wider gender participation disparity than, for instance, the one found in in high-school outdoor track & field, which in 2019 was approximately 55% men to 45% women (“National Federation of State High School Associations” 2019). A smaller participation disparity, like that found in outdoor track & field, should result in a smaller PG due to the relatively higher proportion female athletes in that sport; however, according to my earlier research, the participation disparity does not appear to be correlated with the PG (Carroll 2019).

### Differing Interpretations of the PG Between Men and Women

This perspective is not the only one regarding the PG and human evolution. Some have argued that a wider PG should be used as the evolutionary metric, because it indicates a level of differential selection between the sexes – one that favors males – at certain movements. Morris, Link, Martin, & Carrier (2020), for example, found that men could produce much more relative force than women during a movement that simulated punching versus overhead pulling. And, using the data set from World Athletics, Lombardo and Deaner (2018b) argue that overhead throwing movements like the javelin show a wider PG than either running or jumping as a result of a differential selection favoring males who were superior throwers. These authors all claim that as a result of an evolutionary pressure favoring punching-capable or throwing-capable males, the PG would be wider in punching and throwing movements.

I disagree with these analyses. It is the norm is for men to greatly outperform women in physical ability, not the exception. Many other sports exhibit similarly wide PG’s to throwing: for the heaviest classes of the Olympic lift, the world-record-holding female currently lifts 68.6% the weight of world-record-holding male (“IWF” 2020a; “IWF” 2020b). This PG is lower than the PG of 73.4% between female and male javelin throwers, although it should be noted that this analysis is confounded by weight difference in throwing sports; women throw a 600g javelin, whereas males throw one that is 800g, and by the fact that these Olympic lifting records only reflect two years of competition due to a restructuring of weight classes by the sport’s governing body. Still, were a wider PG linked to differential selection that favored better male throwers, we would not expect such a wide gap between male and female lifters. This example, while certainly limited, does provide more evidence that a wide PG between males and females is the norm and that a narrow PG like that found in rock climbing is more noteworthy.

### Potential Traits That Facilitate Female Climbing

A number of traits likely developed to better facilitate arboreal locomotion; these traits are ones that could close the PG between male and female rock climbers. Firstly, men have a higher bone mass than women (Nieves et al. 2005), which adds additional weight that a man must overcome to ascend a rock face. Secondly, it has been well documented that women on average are more flexible than men, with a greater degree of joint mobility doing movement (Ferber, Davis, & Williams 2003; Kato et al. 2005; Mendiguchia, Ford, Quatman, Alentorn-Geli, & Hewett 2011; Rene 1984). Rock climbing is likely better enabled by increased flexibility, and more-capable rock climbers have been found to possess greater flexibility metrics than less-capable ones (Draper, Brent, Hodgson, & Blackwell 2009; Grant, Hynes, Whittaker, & Aitchison 1996). It should be noted that some studies, however, have called into question the degree to which flexibility impacts climbing performance (Mermier, Janot, Parker, & Swan 2000; Wall, Starek, Fleck, & Byrnes 2004). Finally, and perhaps most importantly, women have been shown to possess greater endurance than men; females are able to maintain isometric contraction for a significantly longer period of time over the same level of relative intensity (Clark, Manini, Thé, Doldo, & Ploutz-Snyder 2003; Hicks, Kent-Braun, & Ditor 2001; Hunter, Critchlow, Shin, & Enoka 2004). This could be in part because women generally have a greater proportion of slower-contractile skeletal muscle fibers, fibers which facilitate endurance rather than power output (Haizlip, Harrison, & Leinwand 2015; Welle, Tawil, & Thornton 2008). Females, though possessing a lower amount of grip strength, have been shown to have levels of forearm and hand endurance equal to or greater than, those of males (Dipla 2017; Fulco et al. 1999; Gonzales & Scheuermann 2007; Hunter 2014). This makes for a climbing advantage in females, muscular endurance being a key component in rock climbing and time to exhaustion a key indicator of a climber’s ability (España-Romero et al. 2009; MacKenzie et al. 2020; Mermier et al. 2000; Giles, Rhodes, & Taunton 2006; Sheel 2004). These traits – lower bone mass, greater flexibility, and better relative endurance – are three physiological examples that could explain in part the extraordinary ability seen in the top female rock climbers.

### Expected Advantages Male Climbers Have Over Females

Interestingly, other traits that make for better climbers are more pronounced in men. A low body fat percentage, for one, is particularly important in climbing (Giles, Rhodes, & Taunton 2006; Watts, Martin, & Durtschi 1993). The body-fat percentage of women is typically higher than that of men, even in elite athletes and rock climbers specifically (Fleck 1983; Mitchell, Bowhay, & Pitts 2011; Novoa-Vignau, Salas-Fraire, Salas-Longoria, Hernández-Suárez, & Menchaca-Pérez 2017). Men, in addition to having greater muscle mass as a percentage of their bodies than women, hold a greater relative percentage of their muscle mass in their upper body versus their lower body (Janssen, Heymsfield, Wang, & Ross 2000). Men also have a cognitive advantage over females in climbing, their heightened visual-spatial processing power (Weiss, Kemmler, Deisenhammer, Fleischhacker, & Delazer 2003). Spatial perception and mental rotation tasks, both of which climbers would rely on during an ascent, are easier for men than for women (Linn & Petersen 1995; Pietsch & Jansen 2018). Taken all together, there do appear to be a number of physiological and cognitive advantages men in general would have over women in climbing, yet women are still adept climbers. This suggests the climbing-specific SBMA’s and sex-blind cognitive adaptations present in humans are numerous and substantial enough to mitigate many of the expected advantages men would have over women in climbing. It also provides evidence that the PG itself may be a better indicator of the evolutionary importance of a movement than individual measures like muscle fiber composition or spatial reasoning.

### How SBMA’s Bridge the Gender Gap in Climbing

A number of attributes are required for successful rock climbing; many of them have been discussed so far in this paper. Some traits, like better flexibility and muscular endurance, are more pronounced in women. Others, like increased upper body muscle mass and heightened visual-spatial reasoning, are more pronounced in men. A third category of traits likely exists, traits that facilitate climbing for both women and for men, and in this category fall the SBMA’s that close the PG between men and women. The structure of the human body – with its many large back muscles, strong hands, and mobile joints – enables tree climbing in a number of ways. These specific adaptations for climbing overcome the general advantage men have at physical activity, a process which manifests itself as a narrower PG between the sexes in the sport of rock climbing. SBMA’s have arisen in response to evolutionary stressors human ancestors faced, like the threat of terrestrial predators, which could explain the narrower PG found in sprinting versus endurance running, for example (Carroll 2019). Although it cannot be ruled out that the narrowness of the gender gap in climbing may in fact be due more to sex-specific traits found in females than to SBMA’s, the extraordinary ability of female climbers relative to other female athletes suggests an evolutionary cause to the PG. While it may be difficult to discern the degree to which any specific trait narrows or widens the PG, the PG itself can likely be used as a measure of accumulated SBMA’s and thus human evolution.

## Conclusion

Arboreal locomotion is well documented as an integral part of human evolution, especially as it relates to primate evolution. As described in this paper, a tremendous volume of evidence suggests that precursors to the genus *Homo*, early *Homo*, and even *Homo sapiens* have all climbed trees; a number of physical and psychological traits found in humans seem at least in part to have originated at least in part to facilitate tree climbing. Many of these traits can be considered SBMA’s, traits common to both men and women, that neutralize the general physical differences between males and females. As a result of SBMA’s bridging the physical gender gap, the PG between men and women narrows in sports that most closely mimic movements essential to human evolution. This is pronounced in rock climbing, an analog of tree climbing, which exhibits a uniquely narrow PG between the sport’s top female and male athletes. The extraordinary ability of female climbers provides additional evidence that a narrow PG is evident in movements essential to human evolution. This information can easily be applied to many movements, allowing researchers to use the PG as another metric with which to evaluate the evolutionary history of our species.

